# Reprogrammable bacterial nanosyringes for RNA and gene editors delivery

**DOI:** 10.1101/2025.10.12.681953

**Authors:** Haishan Xu, Lei Feng, Ningning Song, Xiangxiang Zhao, Xianchao Feng

## Abstract

The engineered *Photorhabdus* virulence cassette (PVC) has not yet achieved the RNA packaging and injection delivery, although it can accurately deliver diverse proteins. Fortunately, RNA-binding proteins have a high affinity for binding to specific sequences and are excellent anchoring components between RNA and the inner tube of PVC. By combining the U1A RNA-binding domain (U1A RBD) and PVC, we developed a convenient, RNA non-degradable, and universal PVC-based RNA delivery tool, establishing **DART (**PVC **D**ocker-based **A**ll-purpose **R**NA Injection Delivery **T**ool). DART successfully achieved the delivery of diverse RNAs, including Pepper RNA, gRNA, siRNA, and miRNA. We also demonstrated that the DART enabled effective knockout of GFP in vivo and in vitro. Our DART system provided a powerful platform for RNA and gene editing therapeutics in vivo.

## Introduction

Delivery technology has emerged as the core engine driving transformative advances in vivo therapeutics^1–3^. By skillfully overcoming physiological barriers (such as cell membrane blockage, enzyme degradation, and immune clearance), it enables the precise delivery of therapeutic nucleic acids (siRNA, miRNA, gRNA), thereby mediating such as gene expression regulation, mutation repair, immune activation, and gene editing in vivo^2,4^. Currently, mainstream carriers can be classified into two categories: virus-derived carriers and nano-derived carriers, which have their own advantages but limitations^5^. Virus-derived carriers benefit from the intrinsic cell-targeting capability of engineered viral capsids. They offer highly precise targeting and prolonged mRNA expression^6,7^, while present limitations including immunogenicity, insertional mutagenesis, and restricted packaging capacity^8^. Nano-derived carriers can be designed to be tailored, are capable of precisely targeting specific sites and mediating transient genome editing. However, their inherent limitations of instability and toxicity pose substantial challenges to clinical applications^9,10^. Accordingly, developing a novel, high-efficiency delivery system that combines the advantages of both virus-derived and nanoparticle-based carriers while minimizing their limitations holds significant applied value.

To overcome these challenges, alternative delivery systems from bacterial origins have recently gained attention. Bacteria-derived *Photorhabdus* virulence cassettes (PVC) is phage-like nanocomplexes capable of targeting eukaryotic cells via a self-injection mechanism^11^. Their unique tubular architecture endows them with an inherent ability to deliver proteins, offering the potential to establish a third, independent delivery paradigm distinct from conventional vectors. PVC can protect their cargo from threats in the cellular environment and can precisely target cells for autonomous injection^11,12^. If they can be loaded with RNA, this will greatly advance RNA and gene editing in vivo. Nevertheless, the narrow inner tube of PVC (length: 1170Å, width: 40Å) necessitates that large cargo proteins adopt an unfolded conformation^11,12^. Thus, selecting a compact protein with high RNA-affinity that is unaffected by spatial constraints is indispensable. Fortunately, U1A precisely meets this requirement, with a simple and stable structure and an RNA-binding domain (RBD) containing only 98 amino acids, which can fit the limited internal tube space of PVC^13^. More importantly, U1A requires only a 9-nucleotide hairpin structure (shRNA) to attain nanomolar-level affinity coupled with high specificity, rendering it an ideal RNA-binding protein^14,15^.

Here, we combined PVC with the U1A RBD to develop a convenient, versatile, and RNA-stabilizing delivery system, termed the PVC **D**ocker-based **A**ll-purpose **R**NA Injection Delivery **T**ool (**DART**). As a proof of concept, different types of RNA including siRNA, miRNA, and gRNA were selected for targeted self-injection delivery both in vitro and in vivo. We delivered gRNA and Cas9 separately and achieved effective co-knockout of GFP in cells and mouse models. Our results demonstrated that the methodology established possesses considerable potential for the delivery of diverse RNAs and gene editors across multiple applications.

## Results

### 1. Strategy design of the DART

The strategy of the DART is illustrated in Figure 1. Briefly, to overcome the limitation of PVC in RNA delivery, among various common RNA-binding proteins such as the MS2 coat protein, PP7, and PUF family proteins^16–18^, we selected the U1A as the RNA anchoring element in the PVC tube^19^. The U1A RNA-binding domain (RBD) not only possesses a simple and stable structure but also exhibits high specificity for binding to the 9-nt loop shRNA^20^. Based on this property, we designed a novel tandem RNA consisting of shRNA and a payload RNA, which can be grafted onto the U1A RBD to form a U1A/payload RNA complex. Under the guidance of the engineered Pd-NTD of U1A, the complex was self-assembled into the inner tube of the PVC (Figure 1a). As designed, we constructed the pPayload RNA plasmid. This plasmid was co-expressed with pPVC in *Escherichia coli* EPI300. DARTs were subsequently purified through ultrahigh-speed differential centrifugation (Figure 1b). During delivery, DART targets recipient cells via receptor recognition and efficiently releases the payload RNA during the puncture process (Figure 1c). Notably, by leveraging the unique targeting and injection mechanism of PVC, this strategy was applied in both in vitro and in vivo settings. It was successfully applied to gene editing, gene silencing, and intracellular imaging, thereby enabling the delivery of diverse RNAs for a broad range of applications^21,22^.

**Figure 1.**
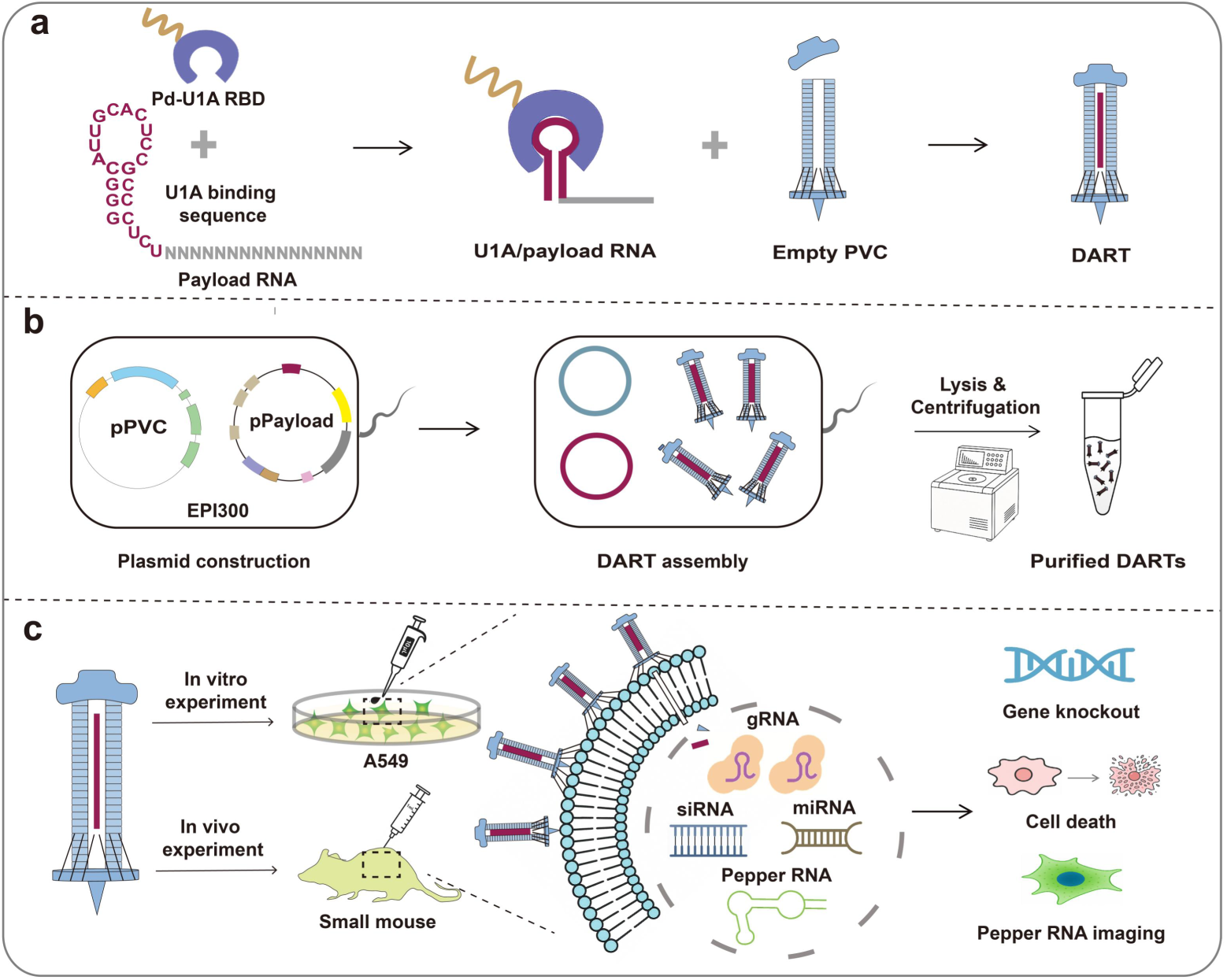
The strategy and work flow of DART. **a** Schematic illustration of payload RNA loads in PVC nanosyringe. Pd-U1A RBD was created by engineering the fusion of U1A RBD (blueviolet) and the leading peptide sequence Pdp1 packaging domain (Pd, golden brown). Tandem RNA contains U1A binding sequence (red purple) and payload RNA (gray). Payload RNA were anchored to U1A RBD by U1A binding sequence, forming U1A/payload RNA complexes. Above these were recruited into empty PVC tubes (sky blue) guided by Pd_NTD, yielding DART, while Pd_NTD itself did not enter into PVC tube. **b** Preparation and purification of DARTs. pPayload and pPVC plasmids were co-expressed in *E. coli* EPI300, and assembled into complexes through regulation. Lysates were processed by ultra-high-speed differential centrifugation to obtain purified DART particles. **c** The application of DART. Purified DART could be applied both in vitro (A549 cells) and in vivo (mouse model). Its self-contractile injection directly delivered diverse RNA cargos—including gRNAs, siRNAs, miRNAs, and Pepper RNAs—into the cytoplasm, enabling CRISPR-based editing, gene interference, and intracellular imaging.

### 2. Validation of DART-based RNA injection in vitro

To validate the feasibility of the RNA injection strategy mediated by the DART, we first designed the Pepper aptamer as the payload RNA (Figure 2a). The Pepper aptamer is an approximately 40-nucleotide RNA molecule that specifically binds to HBC530 to activates its green fluorescence, making it a powerful tool for RNA imaging in live cells and in vitro systems^23–25^. Through the Pepper aptamer, we were able to visually detect whether RNA was loaded into the inner tube of the PVC via U1A and whether it was be delivered into cells (Figure 2a). We constructed the expression plasmid encoding the U1A/payload Pepper complex and subsequently expressed and purified the DART. The related DART-loaded proteins and Pepper RNA sequences were described in Supplementary Table 1. Subsequently, we firstly confirmed the successful loading of U1A into PVC tube through HiBiT detection experiment (Figure 2b) and SDS-PAGE (Figure 2c). Further, the loading of Pepper RNA into DART were verified. The SDS-PAGE and TEM results indicated that DART presented a typical phage-like protein injection structure (Figure 2c-d), containing complete substrate and sheath structures, which suggested that DART was successfully expressed. Furthermore, extracellular fluorescence assays demonstrated that detectable signals were observed only when both Pepper RNA and HBC530 dye were simultaneously present in DART (Figure 2e and Supplementary Figure 1a), with the fluorescence intensity displaying concentration dependence (Figure 2f). These results confirmed that the construction strategy of DART was feasible and that RNA could be successfully loaded into the inner tube of PVC.

**Figure. 2.**
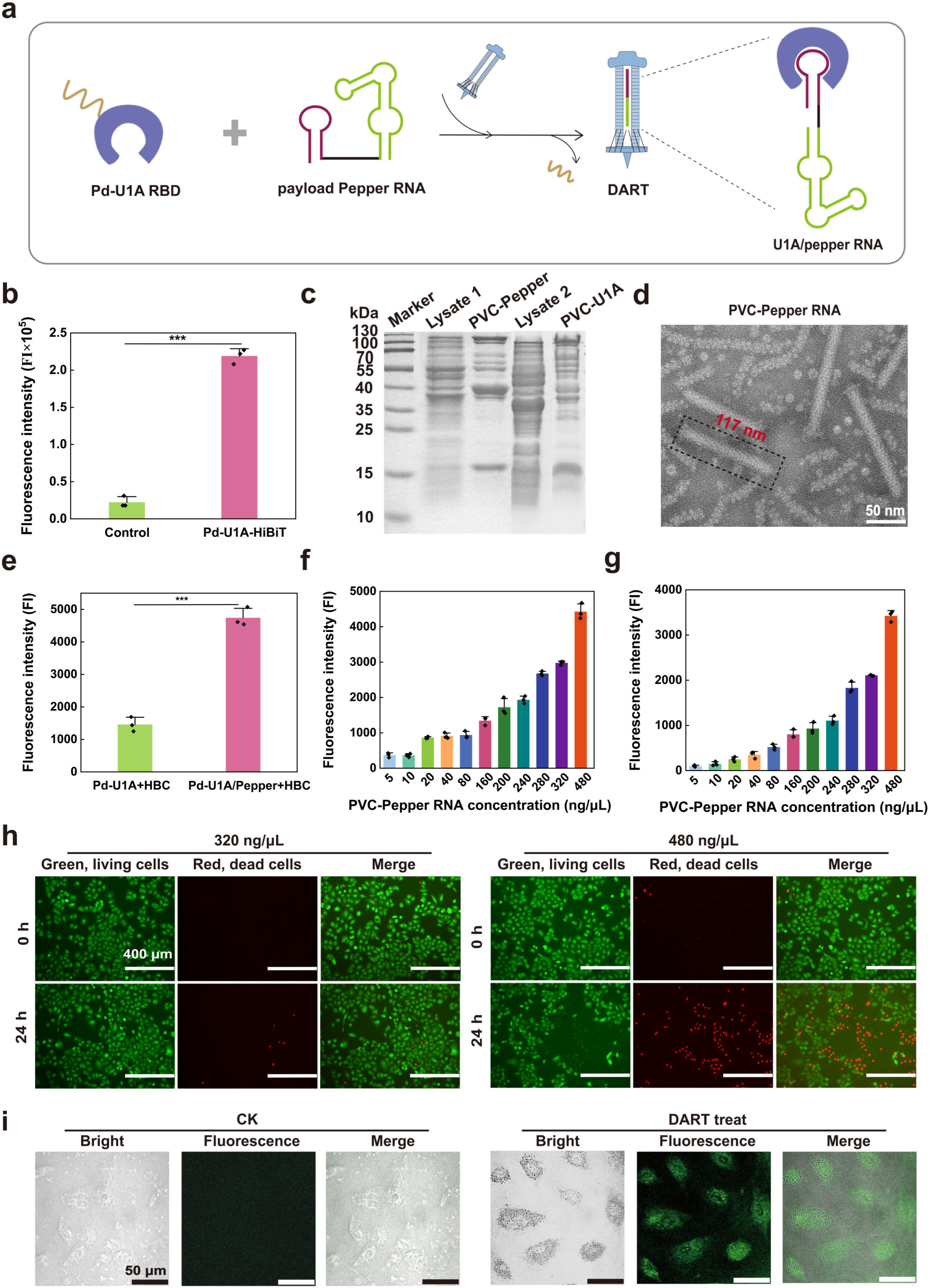
Validation of DART assembly and functional delivery for Pepper RNA. **a** Schematic of DART assembly. The Pepper RNA (green) was fused to the U1A-binding motif (red purple) and recruited into PVC tubes via Pd-U1A RBD. **b** Whether the U1A RBD was successfully loaded was confirmed by HiBiT tag detection. The Pd-U1A-HiBiT group showed a significant fluorescence difference compared with the control group. Data are mean ± SD; n = 3. **c** SDS-PAGE analysis confirmed that U1A and U1A-Pepper RNA were loaded in PVC. **d** Negative-stain TEM images showed phage-tail-like morphology of the DART with an intact sheath and tube (scale bar, 50 nm). **e** Whether the Pepper RNA was successfully loaded into PVC was confirmed by fluorogenic detection coupled with HBC fluorophores. The Pd-U1A/Pepper RNA group showed a significant fluorescence difference compared with the control group. Data are mean ± SD; n = 3. **f** The fluorescence intensity of different concentrations of the DART further confirmed that Pepper RNA was loaded. **g** The cells incubated with the DART showed fluorescence. The fluorescence intensity gradually increased with the increase of DART concentration, which proved that Pepper was delivered into the cells. **h** Live-dead assay confirmed that the optimal concentration of the DART was 320 ng/μL (scale bar, 400 µm). **i** The confocal images of A549 cells showed strong cytoplasmic fluorescence signals after treatment with the DART (right), compared with the control (CK, left) (scale bar, 50 µm).

To evaluate whether the DRAT could efficiently deliver RNA into cells, DART was co-incubated with A549 cells for 2 h, followed by fluorescence analysis of the harvested cells. The results showed that the fluorescence intensity increased with the increasing complex concentration, especially reaching a stable and detectable level at 480 ng/μL (Figure 2g). This confirmed the feasibility of intracellular RNA delivery based on DART. However, when DART concentration exceeded 320 ng/μL, although both intracellular and extracellular fluorescence continued to rise due to the increased RNA loaded (Figure 2 f-g and Supplementary Figure 1b-c), compared with that cells grew well (Figure 2h, left), incubation of A549 cells with high-concentration DART for 24 h resulted in obvious cell death (Figure 2h, right and Supplementary Figure 1d). This finding indicated that elevated DART concentrations increased potential cytotoxicity, highlighting the importance of selecting an appropriate therapeutic dosage. Therefore, considering both delivery efficiency and potential cytotoxicity, we determined 320 ng/μL as the subsequent working concentration of DART. Furthermore, since the co-incubated result could not excludet the possibility of RNA adhering ton the cell membrane, we employed confocal microscopy to perform a detailed intracellular delivery analysis of the DART. The results demonstrated that, compared with the control group (Figure 2i, left), the cytoplasm of cells treated with the DART exhibited distinct green fluorescence (Figure 2i, right). This finding effectively confirmed that the DART successfully delivered RNA into cells. Remarkably, the DART strategy firstly demonstrated stable RNA loading within the enclosed inner tube, making a critical technological breakthrough in achieving intracellular RNA delivery via PVC.

### 3. The versatility of PVC-RNA syringe with double RNA tandem structure

Theoretically, as long as the space of the PVC tube is sufficient, different types of RNAs can be sequentially extended and linked at the 3′ end, allowing the potential co-incorporation of multiple RNAs (Figure 3a). To demonstrate the versatility of RNA loading and delivery by the DART, we extended the downstream region of Pepper with additional RNAs, including siPLK1, miR96 and gRNA^26–28^. The related RNA sequences were described in Supplementary Table 1. Subsequently, the SDS-PAGE and TEM results indicated that no matter which of the above three RNAs were extended to the 3 ‘end of Pepper, they could be loaded into DART and be effectively expressed to form a complete nanosyringe structure (Figure 3b-e). Extracellular fluorescence analysis revealed that, upon incubation of the DARTs with HBC530 dye, fluorescence intensity increased proportionally with DARTs concentration, with pronounced enhancement observed at 480 ng/μL (Figure 3f-h). To assess whether the DART could deliver different types of RNA into cells, we examined intracellular fluorescence following co-incubation of the DART with A549 cells. The results showed that the fluorescence intensity also exhibited concentration dependence (Figure 3i-k). This finding confirmed the generalizability of DARTs for delivering RNA cargo. Although a high concentration of DART allowed more RNAs to be loaded into the tube, it consequently induced strong green fluorescence both intracellularly and extracellularly (Supplementary Figure 1e-j). However, compared at 320 ng/μL (Figure 3l, left), when the DART concentration reached 480 ng/μL or higher, obvious cell death occurred after 24 h-injection (Figure 3l, right and Supplementary Figure 1k). Combing the results of Figure 2, the subsequent working concentration of DART was determined at 320 ng/μL. To further confirm the intracellular fluorescence distribution of DART delivered RNA, we selected A549 cells delivered with gRNA for laser confocal detection and found that distinct green fluorescence was distributed in the cytoplasm (Figure 3m), which further confirmed that gRNA was injected into the cells. In conclusion, the DART enabled stable loading and intracellular delivery of diverse RNA types, offering a universal platform for RNA drug delivery.

**Figure. 3.**
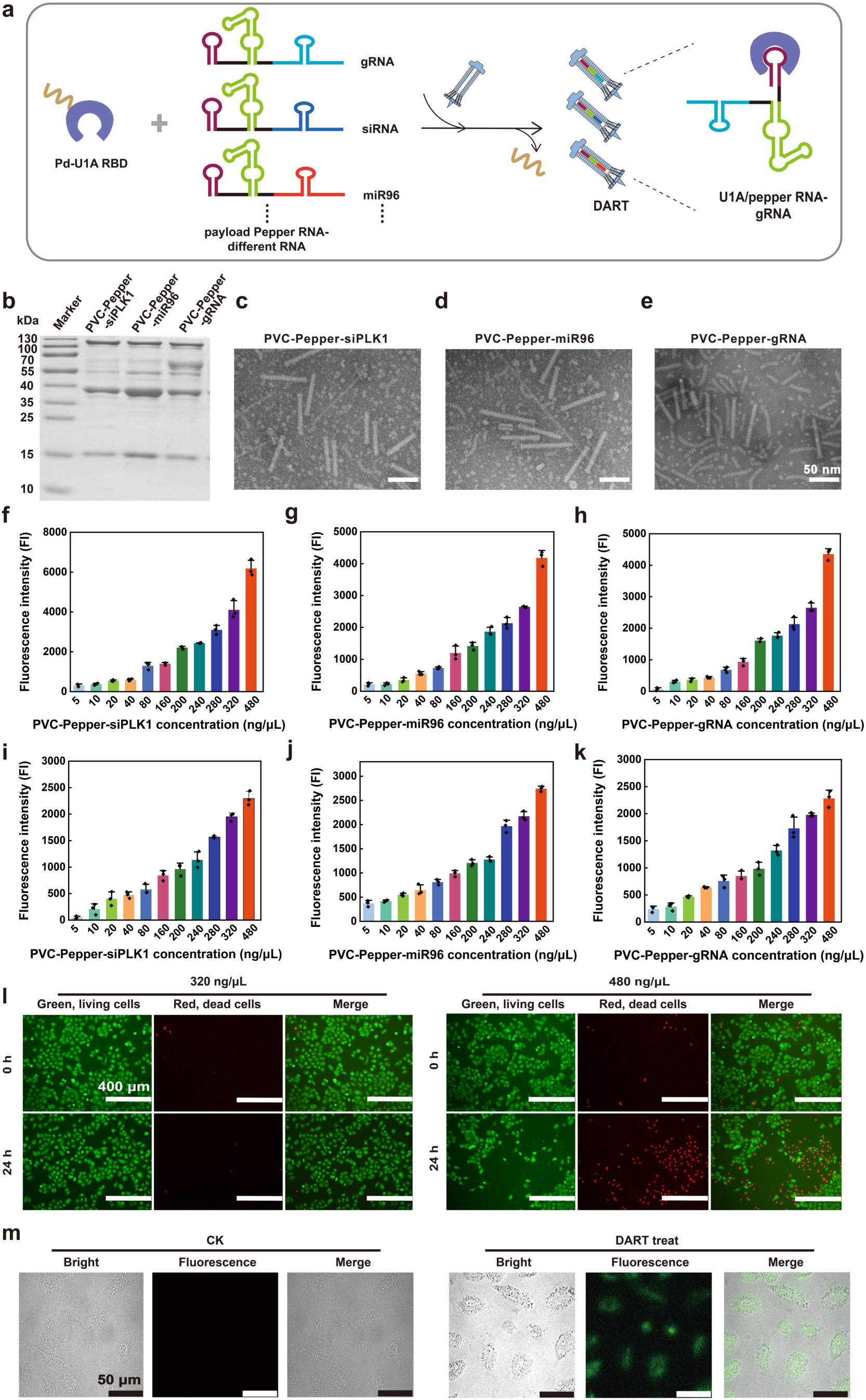
Assembly, characterization, and cellular delivery of different DARTs. **a** Schematic of various DARTs assembly. Payload RNAs (siPLK1, miR96, and gRNA) fused with the U1A-binding motif (red-purple) were anchored to Pd-U1A RBD (blue-violet) and loaded into PVC tubes (sky blue), forming RNA-loaded DARTs, while Pd_NTD itself did not enter into PVC tube. **b** SDS-PAGE analysis proved that DARTs loaded with 3 types of RNAs were successful expression. **c–e** TEM images of DARTs loaded with siPLK1 (c), miR96 (d), and gRNA (e) showed typical phage-like tail morphologies (Scale bar: 50 nm). **f–h** The fluorescence intensity of different concentrations of DARTs further confirmed that siPLK1 (f), miR96 (g) and gRNA (h) were loaded. **i–k** The cells incubated with the 3 types of DARTs showed fluorescence. The fluorescence intensity gradually increased with the increase of DARTs concentration, which proved that siPLK1 (i), miR96 (j), and gRNA (k) were delivered into the cells. **l** Live-dead assay confirmed that the optimal concentration of the DART was also 320 ng/μL (Scale bar: 400 μm). **m** Confocal imaging of DART loaded with Pepper-gRNA in A549 cells, confirmed cytoplasmic delivery (Scale bar: 50 μm). Data are mean ± SD; n = 3.

### 4. Synergistic delivery of CRISPR components achieves gene knockout

Natural PVC is inherently restricted to loading and injecting Cas nucleases^21^. Building on the PVC-RNA syringe, we adopted a split-loading and co-delivery strategy for delivering the CRISPR-Cas system. Specifically, we constructed PVCs individually loaded with Cas9 or Cas12a protein and DART loaded with sgRNA. The plasmid map and agarose gel electrophoresis results were shown in Supplementary Figure 2a-b. These components were then co-added into the same target cells, where intracellular assembly generated functional RNP complexes. The resulting CRISPR RNPs mediated efficient knockout with the intrinsic advantage of transient cleavage.

To verify the feasibility of the co-delivery strategy, we employed the A549-GFP cell line which stably expresses GFP protein as a model system, and designed gRNAs targeting three loci within GFP gene for knockout, including gRNA-G5 and gRNA-G7 corresponding to Cas9 and crRNA-G1 corresponding to Cas12a. The specific gRNA sequences were described in Supplementary Table 4. Fluorescence microscopy revealed that GFP fluorescence intensity gradually increased over 0–96 h in the untreated group and in the group receiving only Cas9 protein or gRNA (Figure 4b). In contrast, in the co-delivery groups (Cas9&G5, Cas9&G7, and Cas12a&G1), GFP fluorescence intensity progressively decreased over 0–96 h (Figure 4b), clearly demonstrating the effectiveness of the co-delivery strategy. Moreover, compared with TatU1A delivery for G5 and PVC deliver for Cas9 (Supplementary Figure 3a), the decrease of GFP fluorescence in the co-delivery groups was more obvious, indicating the better effect of knockout. We further analyzed the cells collected from different treatment groups at 96 h by flow cytometry. The results indicated a similar experimental phenomenon, and the proportion of GFP-negative cells increased (Figure 4c-i). Thereafter, compared with the fluorescence attenuation of different treatment groups, we found that the fluorescence quenching effect were remarkable in the groups of Cas9&G5, Cas9&G7, and Cas12a&G1 (Figure 4g-i).

**Figure. 4.**
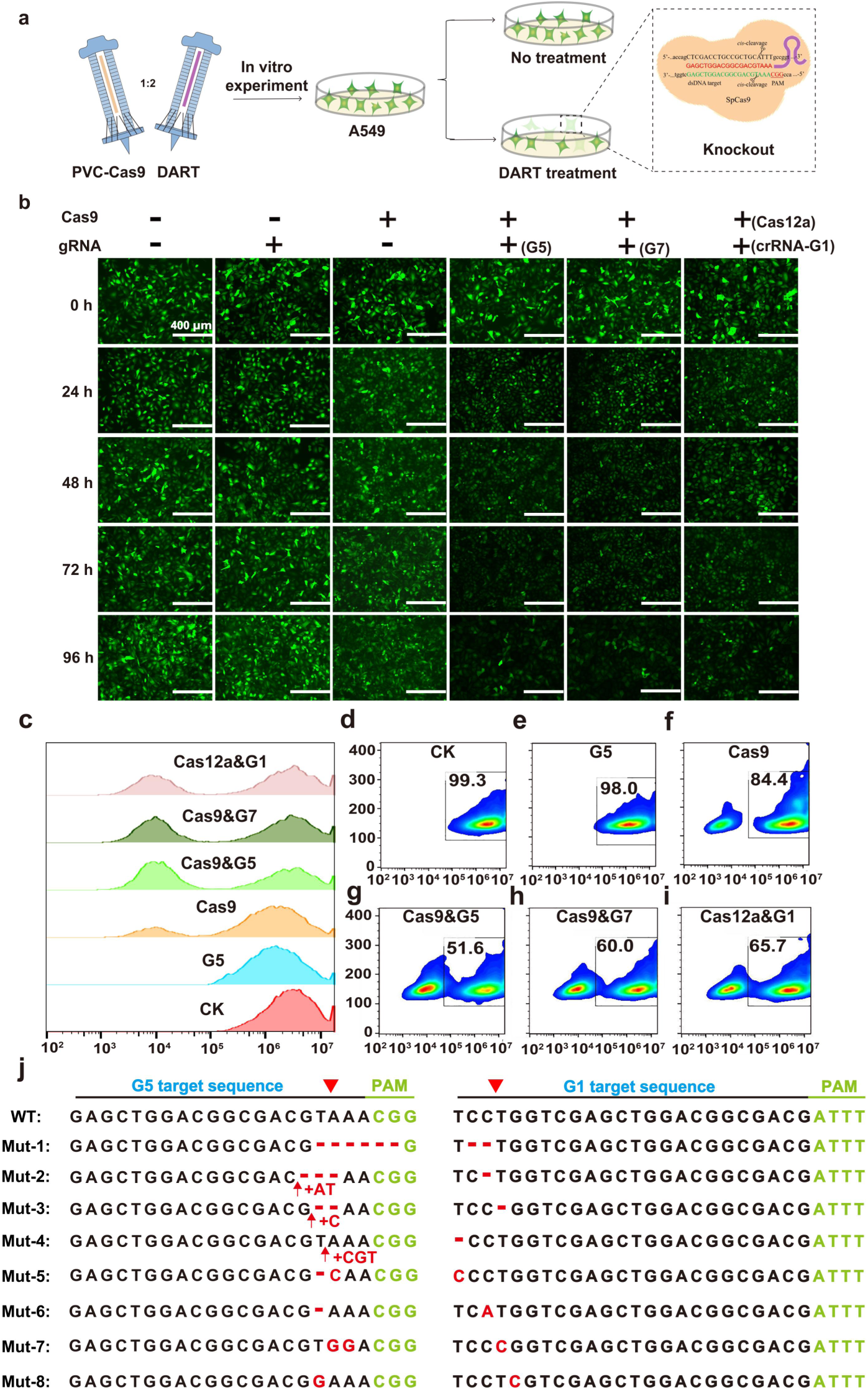
DARTs mediated the knockout of GFP gene in vitro. **a** Schematic of split-loading and co-delivery strategy. PVC particles were separately loaded with gRNAs and Cas proteins (Cas9 or Cas12a), which assembled into active CRISPR RNPs upon delivery into A549 cells. **b** Fluorescence microscopy of A549-GFP cells treated with DARTs. Compared with the control groups (blank control, gRNA, Cas9), the fluorescence of cells in the co-delivery groups (Cas9&G5, Cas9&G7, Cas12a&G1) were significantly quenched. **c–i** Flow cytometry analysis confirmed that co-delivery markedly increased GFP-negative cell populations compared to single-component or control treatments. **j** High-throughput sequencing of target loci revealed indel mutations—including deletions, insertions, and substitutions—following intracellular delivery of Cas9-G5 or Cas12a-G1 via DARTs, confirming efficient genome editing.

To further validate genome editing at the molecular level, genomic DNA was extracted from the treated cells, and specific primers (Supplementary table 5) were designed to amplify the G5 and G1 regions of the GFP locus, followed by high-throughput sequencing analysis. The sequencing results showed that significant indel mutations occurred at all the above-mentioned targets (Figure 4j and Supplementary Figure 3b). The mutation types included single-base insertions, small fragment deletions, and complex base mismatches. The base insertions or deletions would cause frameshift mutations in the GFP gene, further leading to inactivation of GFP protein expression, which was highly consistent with the observed fluorescence attenuation phenomenon.

Subsequently, we further conducted experimental optimization on the knockout efficiency at the G5 and G7 targets. The knockout efficiency of fluorescent genes in cells was significantly enhanced through short-term (6 h) and repeated (12 times) administration. In terms of time, the original action time needed to reach 96 h to achieve a better effect, but the optimized time was shortened to 72 h, achieving a better knockout effect, and the fluorescence intensity of the fluorescent cells decreased significantly (Figure 5a). Further high-throughput sequencing results indicated that base deletions occurred at both the G5 and G7 targets (Supplementary Figure 4a-b), which lead to the inactivation of GFP protein function. Particularly, the knockout at the G5 target increased the proportion of modified reads to 11.58%, resulting in varying degrees of base mutations, substitutions, or deletions (Figure 5b). This further indicates that the G5 target is a favorable knockout site.

**Figure. 5.**
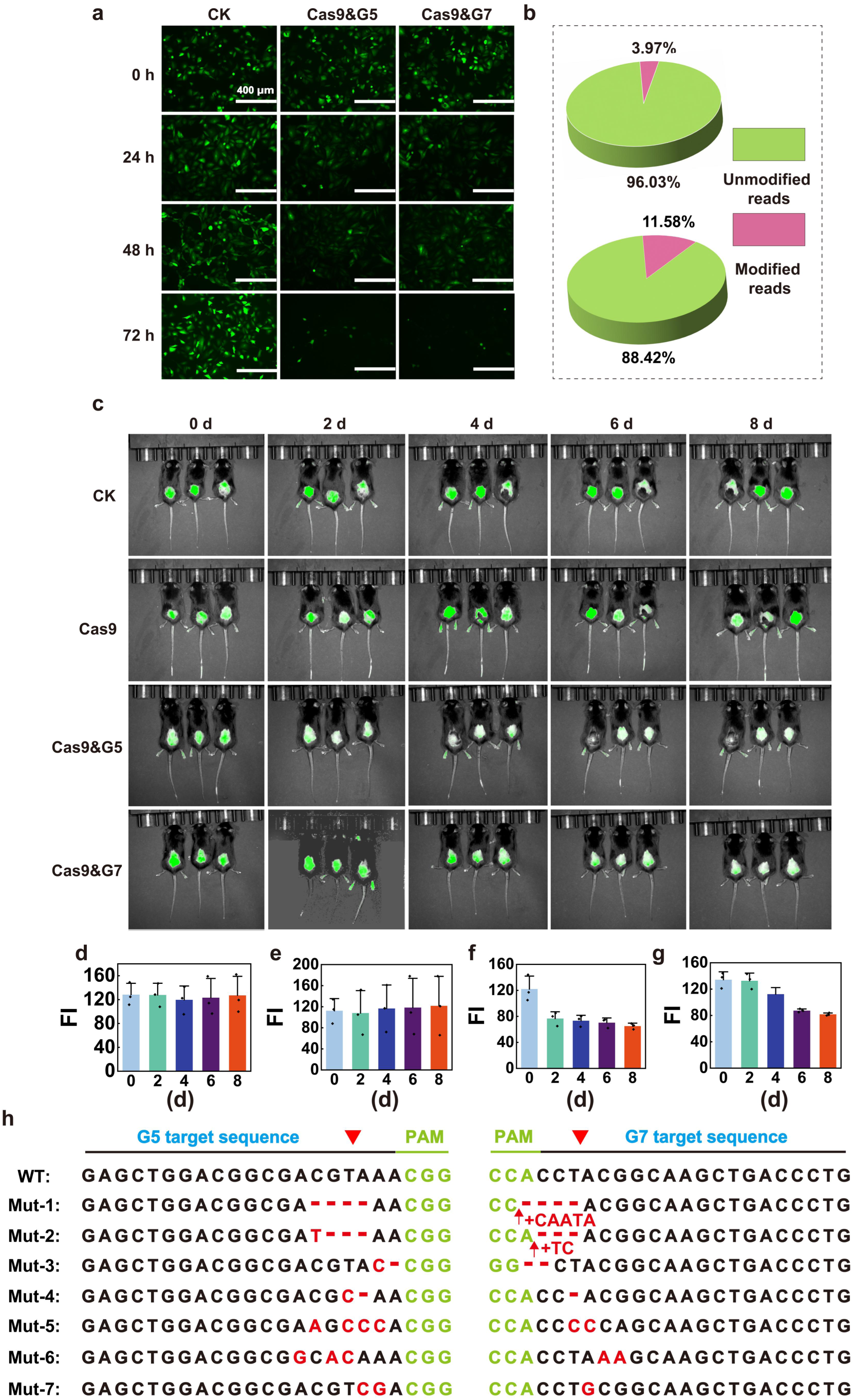
DARTs mediated the knockout of GFP in vivo. **a** GFP fluorescence microscopy of cells after repeated and low-dose co-delivery of Cas9-G5 and Cas9-G7 via PVC particles, imaged at 0, 24, 48, and 72 h post-treatment. Scale bar: 400 μm. **b** High-throughput sequencing quantification of genome editing efficiency at GFP-G5 (top) and GFP-G7 (bottom) target sites, showing the fraction of modified reads. **c** In vivo gene editing in mouse skin. Subcutaneous injection of PVC particles carrying Cas9 and gRNA led to progressive GFP signal loss over 8 days, whereas control groups (untreated or Cas9 only) retained stable fluorescence. **d–g** The fluorescence intensity of mouse skin measure within a fixed region of interest using image. d represents CK, e represents Cas9, f represents Cas9&G5 and g represents Cas9&G7. **J** It also indicated the extensive GFP signal loss under the above same treatments. **h** Representative indel patterns at GFP-G5 (left) and GFP-G7 (right) target sites revealed insertions, deletions, and substitutions, many causing frameshift mutations.

In order to further verify the gene knockout capability of DART in living organisms, DARTs co-loaded with Cas9 and G5/G7 were delivered to mouse skin tissue by subcutaneous injection. Fluorescence imaging was performed every two days following injection. The results showed that the green fluorescence signal in the skin tissue of the control group remained stable throughout the observation period, whereas in the co-delivery group, the fluorescence gradually declined with the increase of time (Figure 5c-g). Especially for Cas9&G5 treated group, the fluorescence significantly decreased from the second dayits and was remarkly reduced by the 8th day (Figure 5d-g). Even in certain regions, the fluorescence signal nearly disappeared, indicating successful activation of the CRISPR system in vivo and effective knockout of the target gene. To further confirm the occurrence of the editing event at the molecular level, genomic DNA of the processed tissue was extracted, PCR amplification was performed on the GFP-G5 target site, and high-throughput sequencing analysis was carried out. The results showed that significant mutations occurred in the target region, including base insertions, small fragment deletions and complex variations (Figure 5h). The above results jointly confirmed that the DART enables precise delivery and functional activation of CRISPR components in vivo, highlighting its broad potential for gene editing, tissue-specific regulation, and disease model generation. Moreover, it was found that DART became hydrolyzed after 8 d treatment skin tissue, indicating DART could be rapidly clearance and was safe (Supplementary Figure 4c).

In conclusion, DART not only facilitates efficient gene editing in cells but also enables precise and safe in vivo delivery via self-contracting injection without the need for complex instruments.

### 5. The widespread and cross-species applicability

Previous studies have demonstrated that PVC can be engineered to recognize specific target cells, thereby conferring precise delivery capabilities^11,21,29^. In this study, we employed AlphaFold3 to predict the structure of the engineered DART tail fibers (Supplementary Figure 4a), facilitating the application of the DART across a broader range of species. We selected the AD5RGDPK7 or DARPin domains,which have been shown to specifically recognize integrin receptors or growth factor receptors, respectively^11^. We performed structural modeling of surface integrin or growth factor receptors across multiple model organisms (including humans, monkeys, pigs, rats, mice, zebrafish, fruit flies and nematodes) (Supplementary Figure 4b-i), and predicted the recognition and binding of the engineered tail fibers to the receptors through computer-aided methods (Figure 6a)^30^. We found that the DART exHiBiTed strong receptor-specific recognition potential, enabling its broad applicability across diverse model organisms, from higher to lower species. In addition to humans and mice, we selected zebrafish, which have a relatively large species gap, for further verification.

**Figure. 6.**
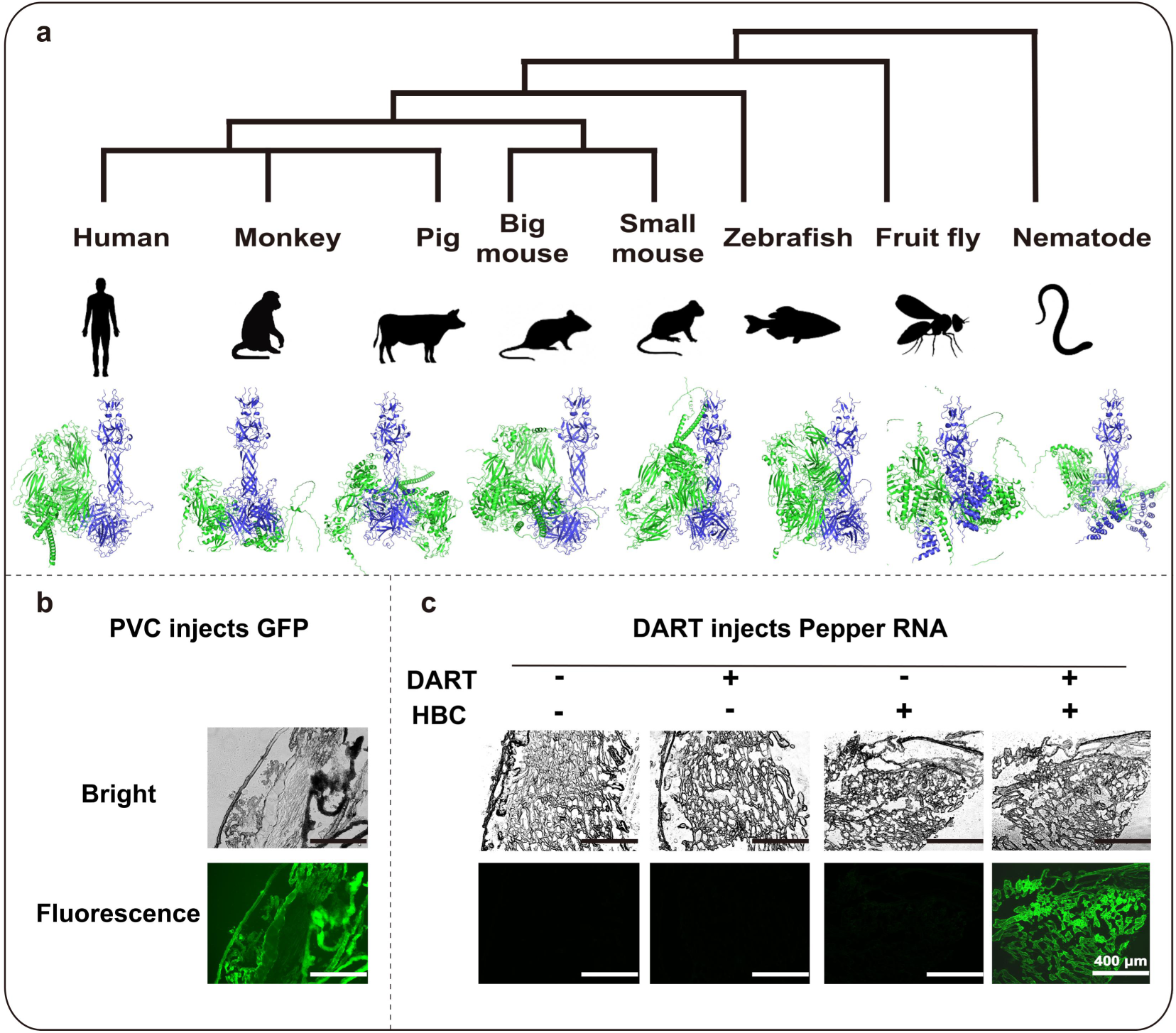
Broad-spectrum applicability of engineered DARTs across species. **a** Phylogenetic distribution of representative model organisms—including human, monkey, pig, rat, mouse, zebrafish, fruit fly, and nematode—used for structural prediction. AlphaFold3.0 modeling of engineered PVC tail fibers fused with AD5RGDPK7 or DARPin domains and corresponding host receptors was performed. **b** Representative GFP delivery in zebrafish skin via PVC injection, showing robust fluorescence, whereas control pnf delivery caused lethality. **c** Validation of Pepper RNA delivery into zebrafish tissues confirmed effective cytoplasmic delivery.

To experimentally validate these predictions, we first injected GFP-loaded PVC particles into zebrafish. Strong green fluorescence was observed (Figure 6b), demonstrating that PVC could specifically recognize zebrafish and delivered its protein cargo. We then injected zebrafish with DART particles loaded with Pepper RNA. Notably, green fluorescence was only detected when both DART and HBC were present (Figure 6c), indicating that DART could specifically recognize zebrafish surface receptors and achieve precise RNA delivery. Together with our previous findings of successful RNA delivery into human and mouse cells, these results further supported the cross-species applicability of the DART platform.

## 6. Discussion

The successful realization of RNA delivery based on PVC is of considerable significance. Compared with protein and small molecule drugs, RNA lies in their high specificity and designability can target any gene and have the potential for rapid development and precise intervention^31,32^. However, RNA itself is unstable, easily degraded by nucleases in the body, and carries a negative charge, making it difficult to cross the cell membrane and enter functional areas within the cell^33^. Therefore, a novel and efficient delivery system is indispensable. However, traditional delivery systems still face many challenges in terms of delivery efficiency, tissue specificity, safety and controllable release^34^.

This study addressed the key bottlenecks in the in vivo delivery of RNA drugs and gene editing systems and provided a bacterial-derived vector different from viral- and nanomaterial-derived vectors^6,8^. By systematically researching bacterial functional components, we upgraded the *Photorhabdus*-derived PVC, and expanded its capabilities. These modifications overcame the original constraints of the limited PVC tube and its ability to load only protein cargo^11^. We developed a convenient, nuclease-resistant, and universally programmable PVC **D**ocker-based **A**ll-purpose **R**NA Injection Delivery **T**ool. This system allows plug-and-play replacement of diverse RNA molecules, thereby enabling efficient RNA loading and delivery within the confined and limited inner tube of PVC. Specifically, we took advantage of the characteristic that U1A can specifically bind to tandem RNA containing shRNA sites and has a simple structure that can adapt to the narrow inner tube of PVC. We introduced U1A RBP as an “anchoring plate” and designed a serial tendam RNA containing shRNA sites to form the U1A/U1A-payload RNA complex^35^. Unlike the recently reported SPEAR system, which fused different cargo (RNP and ssDNA) onto the spike complex proteins Pvc8/Pvc10 and mediates loading and delivery outside the spike^21^. Our work introduced the small-sized and highly affine U1A RBP at the PVC inner tube, achieving stable loading of RNA within the inner tube. Consequently, DART represents another significant innovation in PVC delivery research. And our strategy can significantly reduce the risk of degradation of exposed RNA and leverage the delivery advantages of PVC self-contractile injection, truly achieving the leap from “protein delivery” to “RNA delivery”. DART has a high degree of modularity and universality. It only needs to replace the 3’ terminal functional region sequence of the tandem RNA to achieve the loading and delivery of different RNAs. In this study, the DART successfully achieved “plug-and-play” loading of multiple RNA types, including siPLK1, miR96 and gRNA, as evidenced by both intracellular and extracellular fluorescence measurements and confocal microscopy analyses. This design implies that DART is no longer limited to a single payload, but can be flexibly combined according to therapeutic needs to support the simultaneous delivery of multiple RNAs. It thus holds the potential to serve as a standardized delivery platform for RNA-based therapeutics.

In terms of functional verification, this study not only demonstrates the efficient loading and delivery of RNA molecules by the DART, but also highlights its potential for gene-editing applications. Specifically, gRNA and Cas9/Cas12a protein were independently loaded into the DART, respectively, and co-delivered to A549 cells to achieve GFP gene knockout, leading to gradual quenching of cellular fluorescence. Not only that, we successfully achieved in vivo GFP gene knockout in a fluorescent transgenic mouse model, demonstrating that the DART can be effectively applied in living organisms. Unlike approaches relying on electroporation, microinjection, the DART can precisely deliver cargo to cells without the need for complex instruments owing to its intrinsic stretchable injection mechanism^36–38^. In the near future, it is also very likely that oral or inhalation administration methods will be realized, which will further enhance the convenience of DART. And the DART production workflow is straightforward and cost-effective. Its cost, as low as $2.3, was 10 to 1,000 times lower than that of the existing delivery vectors (AAV, Lentivirus, etc.) (Supplementary Table 6 and 7). Additionally, the DART exHiBiTs excellent RNA protection capabilities, which provides a new type of bacterial source delivery tool independent of viral and nanomaterial source vectors for the delivery of RNA drugs and gene editing systems.

In conclusion, this study established the first DART with programmable and modular features, thereby extending PVC to the RNA delivery level. Once the expression plasmid is constructed, it can be used to produce syringes once and for all, which is not only convenient but also cost-effective. Compared with SPEAR’s approach of pushing PVC towards the external surface engineering of “protein /RNP/ssDNA” through “spike-external fusion-in vitro reconstruction”, this study pushes PVC towards the internal engineering, thus providing a parallel and complementary engineering route to SPEAR.

## Materials and Methods

### Plasmid construction

Plasmids encoding PVC structural genes (pAWP78-PVCpnf_pvc13-Ad5RGDPK7 and pAWP78-PVCpnf_pvc13-E01DARPin) and regulatory elements (pBR322-Pdp1_NTD-GFP-HiBiT) were obtained from Addgene (IDs:https://www.addgene.org/popular-plasmids/). The gene of Cas12a was copied from pET28TEV-LbCpf1 which is stored in our lab^39,40^. All PCR amplifications for cargo and regulatory plasmid (pPayload) assembly used Phusion Flash 2× Master Mix (Thermo Fisher Scientific, F548S). Constructs were transformed into chemocompetent *E. coli* DH5α via T5 exonuclease-dependent assembly (TEDA; Laboratory-prepared). EPI300 cells (Laboratory stock) harboring PVC structural plasmids (pAWP78-PVCpnf_pvc13-E01DARPin or pAWP78-PVCpnf_pvc13-Ad5RGDPK7) were electroporated with pPayload. Oligonucleotide synthesis and Sanger sequencing were performed by General Biotech (China). Plasmid inventories and Prime sequence data were cataloged in Supplementary Tables 2 and 5, respectively.

### In vitro transcription of RNA

RNA was transcribed in vitro using the MEGAscript™ T7 Kit (Thermo Fisher, AM1334). Linearized DNA templates (1 μg) with T7 promoters were incubated with T7 RNA polymerase, 1× reaction buffer, and 75 mM NTPs at 37°C for 16 h. Reactions were treated with Turbo DNase (1 U/μg DNA, 37°C, 1 h), purified using the MEGAclear™ Kit (Thermo Fisher Scientific, AM1908), resuspended in RNase-free water, and stored at −80°C.

### DART expression and purification

DART expression and purification followed the method of Kreitz et al^11^. with minor modifications. Single colonies were inoculated into LB medium with kanamycin (50 μg/mL) and ampicillin (100 μg/mL), grown at 37°C (200 rpm, 12 h), then diluted 1:1000 into 500 mL fresh LB with antibiotics and induced at 30°C (160 rpm, 24 h). Cell pellets were resuspended in lysis buffer (25 mM Tris-HCl pH 7.5, 140 mM NaCl, 3 mM KCl, 5 mM MgCl₂) supplemented with protease inHiBiTor cocktail (1×), DNase I (25 mg/mL), and lysozyme (100 μg/mL), then incubated at 25°C for 90 min. Lysates were clarified by centrifugation (4,000 ×g, 30 min, 4°C). Supernatants were ultracentrifuged (120,000 ×g, 2 h, 4°C), and pellets were resuspended in PBS for secondary purification (16,000 ×g, 15 min; 120,000 ×g, 2 h). To get pure DART, the resulting supernatant was centrifuged (120,000 ×g, 2 h, 4°C). After resuspending it in 300 μL of sterile PBS, use a Implen NanoPhotometer N80 (Implen, Germany) to measure the DART concentration based on the A280 value. Note: The purified DART was kept at 4°C in sterile PBS, and it is advisable to utilize it within 7 days.

## Characterisation of DART structures

### Verification of DART protein structure

Purified DART samples were denatured in 6× SDS loading buffer (375 mM Tris-HCl pH 6.8, 12% SDS, 60% glycerol, 0.6 M *β*-mercaptoethanol, 0.06% bromophenol blue) at 95°C for 10 min, resolved on 4%/12% discontinuous polyacrylamide gels, and stained with Coomassie Brilliant Blue R-250 (Thermo Fisher Scientific, LC6025, 40 % methanol, 10 % glacial acetic acid) and stained at room temperature for 30 min on a shaker (Changzhou Nuoji Instrument Co., Ltd, China). After staining, the dye solution was discarded and the gels were subjected to destaining with a solution containing 50% (v/v) methanol, 10% (v/v) glacial acetic acid, and 40% (v/v) double-distilled water on a shaker. The destaining solution was refreshed several times until the background became clear and protein bands were well resolved. Images of the destained gels were captured under visible light using ChemiDoc™ MP Imaging System (BIO-RAD, USA)^41^.

### Microscopic morphology observation of DART

In order to explore the potential effect of RNA loading on the microscopic morphology of DART particles, negative-staining TEM analysis was performed by Scientific Compass (Beijing, China), an ISO/IEC 17025-certified independent analytical laboratory. The precise steps are as follows: DART samples (100–500 ng/μL) were adsorbed onto plasma-cleaned 200-mesh carbon-coated copper grids, negatively stained with 2% uranyl acetate, and imaged using a JEOL JEM-2100F TEM at 200 kV (Gatan US1000 CCD camera, low-dose mode). Images of representative visual fields are used for subsequent morphological analysis^42^.

### Fluorescence detection of DART

Fluorescence detection of DART loaded with Pepper RNA was performed to evaluate the specificity and efficiency of DART in generating fluorescence through the HBC530–Pepper RNA interaction. RNA delivery efficiency was quantified via the Pepper RNA–HBC530 fluorogenic system^43^. Samples were incubated with HBC530 (180 nM final) for 15 min in the dark. Fluorescence (ex/em: 480/520 nm) was measured using a SpectraMax i5x plate reader (Molecular Devices, USA). In addition, PVC loading of U1A protein was verified using the Nano-Glo® HiBiT Extracellular Detection System kit (Promega, NG2421), with all procedures performed according to the manufacturer’s instructions. Net fluorescence intensity (Net FI) was calculated by subtracting reagent and sample blanks. The result was then recorded as Net FI. Note: To prevent experimental contingencies, three technical replicates were assigned to every sample.

#### Cell culture

The human non-small cell lung cancer cell line A549 and its derivative stably expressing firefly luciferase and green fluorescent protein (OriCell-A549-Luciferase-GFP) were obtained from Cyagen Biosciences (China) (Supplementary Table 3). Cells were cultured under standard conditions in T25 flasks (Corning) at 37°C with 5% CO₂ and saturated humidity in a thermostatic incubator (Thermo Scientific Heracell™ 150i, HERAcell 150i, USA) (Kreitz et al., 2023). The basal medium was DMEM/F12 (1:1) (Purino, PM150312), supplemented with 10% (v/v) Gibco heat-inactivated fetal bovine serum (Thermo Fisher Scientific, 26010074) and 1% (v/v) penicillin–streptomycin (10,000 U/mL penicillin, 10,000 μg/mL streptomycin; Thermo Fisher Scientific, 15140122). Cells were routinely passaged using 0.25% Trypsin–EDTA (Thermo Fisher Scientific, 25200056).

### Fluorescence detection of DART intracellular delivery RNA

To evaluate RNA delivery efficiency mediated by DART, we applied a fluorescence-based assay exploiting the Pepper–HBC interaction. A549 cells (Fisher scientific, 50672592) were seeded in black, clear-bottom 96-well plates (VWR 89091-012) at a density of 1 × 10⁴ cells per well and maintained in DMEM/F12 medium supplemented with 10% FBS at 37°C in a humidified 5% CO₂ incubator. Once cultures reached approximately 80 ± 5% confluence, the medium was exchanged for 90 μL of fresh DMEM/F12. DART carrying the target RNA was then introduced to achieve final RNA concentrations of 0, 5, 10, 20, 40, 80, 160, 200, 240, 360, and 480 ng/μL, with the total volume adjusted to 100 μL. Each concentration was assayed in triplicate. After 3 h incubation, cells were harvested, pelleted at 12,000 ×g for 10 min at 4°C, and resuspended in 20 μL of ddH₂O. Suspensions were transferred into black 96-well plates, combined with 4 μL of 2 μM HBC530 (final concentration ∼180 nM), and incubated for 15 min in the dark. Fluorescence was recorded on a microplate reader (SpectraMax i5x, Molecular Devices, USA) at 480 nm excitation and 520 nm emission^44^.

### Laser confocal detection of intracellular delivery of RNA using DART

To visualize intracellular RNA delivery mediated by DART, real-time imaging was performed using the Pepper–HBC fluorescence system. A549 cells were seeded in 29 mm glass-bottom dishes at a density of 8 × 10⁴ cells per well. Upon reaching ∼80% confluence, DART loaded with Pepper RNA–gRNA were added to the culture medium at a final concentration of 3.6 μg/μL. After 3 h incubation, the medium was removed and cells were gently rinsed three times with prewarmed PBS (pH 7.4) to eliminate residual DART. Cells were then fixed in 4% paraformaldehyde (Beyotime, P0099-100ml) for 15 min at room temperature, followed by three PBS washes to remove fixative. Fluorescence imaging (excitation 480 nm, emission 520 nm, Leica TCS SP8, Germany) was performed with a laser scanning confocal microscope, and representative images were acquired for analysis^45^.

### A549-EGFP fluorescence gene knockout experiment under DART treatment

Single-guide RNAs (sgRNAs) targeting the enhanced green fluorescent protein (EGFP) gene were designed using online tool (https://zlab.squarespace.com/ guide-design-resources). Candidate target sites were selected as 20-nucleotide sequences located immediately upstream of the PAM motif (NGG) or 23-nucleotide sequences immediately upstream of the PAM motif (TTTN). Three gRNAs with high scores and specificity (gRNA-G5, gRNA-G7, and gRNA-G1) were selected for further analysis. Plasmid construction procedures followed the methods described in the “Plasmid construction” section. The plasmid inventories and gRNA sequence data are cataloged in Supplementary Tables 2 and 4, respectively.

The dual-reporter A549 cell line (Purino Biological Co., Ltd. (China)) was employed to assess gene knockout mediated by DART. Control groups comprised untreated cells, gRNA-only cells, and Cas9-only cells, whereas experimental groups included Cas9&G5-, Cas9&G7-, and Cas12a&G1-treated cells. Each condition was tested with six biological replicates, and the entire experiment was independently repeated three times. For assays, A549 cells were seeded at 1 × 10⁴ cells per well in clear-bottom 96-well plates and allowed to attach for 12 h. Thereafter, DART loaded with RNA and PVC loaded with Cas9 were mixed at a concentration ratio of 1:2 to prepare the stock solution system, which was then added to 96-well plates to a final concentration of 320 ng/μL. Culture medium was refreshed every 24 h, with PVC complexes re-supplemented at the same concentrations, until cells were collected at 96 h for genome-editing analysis.

### Fluorescence detection of A549-EGFP cells under DART treatment

Fluorescence images were acquired at 0, 24, 48, 72, and 96 h using a fluorescence microscope (scale bar, 400 μm, NIKON, Ti2-U, China) to monitor temporal changes in cell fluorescence intensity and assess the gene-editing activity of the DART-delivered CRISPR–Cas system.

Flow cytometry was performed at 0 and 96 h to quantify fluorescence changes and evaluate the gene-editing effect of the DART-delivered CRISPR–Cas system on the EGFP reporter. Cells from each treatment group at 96 h were harvested, centrifuged, and resuspended in PBS at 1 × 10⁶ cells/mL. Samples were filtered through a 40 μm mesh prior to analysis on a CytoFLEX flow cytometer (Beckman Coulter life sciences, USA). GFP was excited at 480 nm, and emission was collected using a 520/30 nm band-pass filter^46^. Debris was excluded based on FSC/SSC gating, and single cells were gated using FSC-A versus FSC-H. For each sample, 1 × 10⁴ events were recorded. Data were analyzed using FlowJo v10.8.1.

### High-throughput sequencing of cells under DART treatment

To quantify gene knockout mediated by DART, cells treated for 96 h were collected, and genomic DNA was extracted using the TransDirect® Animal Tissue PCR Kit (TransGen Biotech, AD201). DNA was used directly as a PCR template to minimize fragmentation. The target locus was amplified with NEBNext High-Fidelity 2× PCR Master Mix, and purified amplicons were subjected to high-throughput sequencing (Nuohe Zhiyuan Biotechnology Co., Ltd., Beijing, China). Sequencing data were analyzed using CRISPResso2 to determine the frequency of deletions/insertions/mutations (indels) and characterize editing event types.

### Gene knockout experiment of in vivo fluorescent mice under DART treatment

C57BL/6-Tg(CAG-EGFP)131Osb/LeySopJ transgenic mice (Shulaibao(Wuhan) Biotechnology, China) were maintained in a specific-pathogen-free (SPF) facility at 22 ± 1 °C and 55 ± 5% humidity under a 12 h light–dark cycle. Animals had ad-libitum access to sterilized 5CJL-LabDiet® JL Rat and Mouse Auto 6F feed (Yongli Biological, 1813001) and purified water filtered through 0.22 μm membranes (Wahaha, China). All procedures were conducted in accordance with AAALAC guidelines and the GB/T 35892-2018 standard.

The gRNAs (gRNA-G5 and gRNA-G7) corresponded to those used in the A549 EGFP gene-editing experiments under DART treatment. Experimental groups comprised Cas9&G5- and Cas9&G7-treated mice, while controls included untreated and Cas9-only groups. For in vivo gene editing, the dorsal hair of mice (3 × 3 cm) was shaved and disinfected with 75% ethanol. PVC complexes loaded with Cas nucleases and DART complexes containing U1A protein–single-stranded guide RNA were subcutaneously injected at final doses of 1 μg and 2 μg per injection, respectively. Injections were repeated every 24 h for a total of four administrations. Following the experimental period, mices were sacrificed according to AAALAC and GB/T 35892-2018 ethical standards, and genomic DNA was extracted from skin tissue using an animal tissue DNA extraction kit (Simgen, 3101050) under BSL-2 laboratory conditions. Subsequent high-throughput sequencing was performed as described in “High-throughput sequencing of cells under DART treatment.”

### GFP and Pepper RNA delivery of in vivo zebrafish under DART treatment

To assess the delivery capability of the DART across various model organisms, zebrafish (Danio rerio) were chosen as the representative experimental subject. Initially, GFP was delivered into zebrafish using PVC to confirm its targeting efficacy in this species. Subsequently, DART loaded with Pepper RNA solution (3.6 µg/µL, 20 µL per tail) was intraperitoneally injected into the zebrafish, with the injection site located approximately 2-3 mm anterior to the colonization orifice. After 3-h injection, zebrafish were dissected under aseptic conditions. Skin tissues were collected for embedding and rapid freezing, and tissue sections were prepared utilizing a cryosectioning machine (CM1860, Leica Microsystems, Wetzlar, Germany). For the GFP delivery group, sections were directly imaged using a fluorescence microscope (scale bar, 400 μm, NIKON, Ti2-U, China). For the Pepper delivery group, HBC530 dye (6 µL at 2 µM) was applied onto the sections and allowed to stain in darkness for 5 min prior to imaging. Three control groups were established: I) Only HBC530 dye was added II) Only Pepper was delivered without HBC530 III) Neither Pepper nor HBC530 was administered.

### Statistics and reproducibility

All data were analyzed using IBM SPSS Statistics 23. Each experimental condition included three independent replicates, and quantitative results are presented as mean ± standard deviation. Parallel data refer to measurements obtained under identical experimental conditions unless otherwise specified. Representative images from gels and microscopy are shown. For multiple comparisons, one-way or two-way analysis of variance with Bonferroni post hoc correction was applied. Statistical significance was as *P* < 0.05 (*), *P* < 0.01 (**), and *P* < 0.001 (***).

## Author statement

**Haishan Xu and Lei Feng**: contributed to Investigation, Methodology, Data curation, Writing – original draft. **Ningning Song**: contributed to Software, Formal analysis and Conceptualization. **Xiangxiang Zhao and Xianchao Feng**: contributed to Supervision, Resources, Funding acquisition, Writing – Review & Editing.

## Acknowledgements

This work was supported by the National Natural Science Foundation of China (grant nos. 32401252, 32172143 and 318732014)), and the Key Research and Development Program of Shaanxi Province (Program No. 2024SF-ZDCYL-03-23). We are deeply grateful to all colleagues and collaborators for their valuable contributions, insightful discussions, and technical assistance throughout this study. We also sincerely acknowledge the broader community whose support facilitated the completion of this work. Finally, we respectfully honor the experimental animals whose sacrifices enabled the progress of this research, and we reaffirm our commitment to the highest standards of animal welfare and ethical research practices.

## Declaration of Competing Interest

The authors declare no competing financial interest.

## Notes

### Competing Interest Statement

The authors have declared no competing interest.

