## Supporting Information for "Reprogrammable bacterial nanosyringes for RNA and gene editors delivery"

#### Table of contents

Research Overview 1

Supplementary Figures 2

Supplementary Figure 1 Fluorescence of DARTs loaded with different RNAs 2

Supplementary Figure 2 The construction of pBR322-Pdp1_NTD-Cas12a 3

Supplementary Figure 3 TatU1A mediated the knockout of GFP gene in vitro 4

Supplementary Figure 4 PVC tail fiber microstructure and receptor-targeting models 5

Supplementary Tables 6

Supplementary Table 1 Design of PVC-loaded proteins and RNAs sequences 6

Supplementary Table 2 Synthesis and preparation of DNA or RNA 8

Supplementary Table 3 Prime sequence 9

Supplementary Table 4 Strains, plasmids, and primers used in this study 11

Supplementary Table 5 Cell lines used in this study 12

Supplementary Table 6 Comparison of DART with existing delivery vectors 13

Supplementary Table 7 Cost calculation of DART (per 1 mL) 14

Supplementary Methods 15

Supplementary References 16

### Research Overview

In this study, a PVC **D**ocker-based **A**ll-purpose **R**NA Injection Delivery **T**ool (**DART**) was innovatively developed. The U1A RBD was integrated into the PVC contraction injection system to achieve modular loading and efficient delivery of a variety of functional RNAs. Using the characteristics of micro U1A RBD with only 98 amino acids to specifically recognize shRNA, a tandem RNA vector was designed to break through the bottleneck that PVC could not load RNA and overcome the problem of inner tube size limiting protein-RNA complex loading. By replaced the 3' end extension sequence of tandem RNA, universal delivery of different RNA types such as siRNA, miRNA, and gRNA were achieved, and the structural integrity of RNA is maintained. The DART delivery Cas protein and gRNA technology was used to successfully knock out GFP in vivo and in vitro. In order to comprehensively support the conclusions of this study and ensure the integrity of the experimental system, the supplementary materials integrate multiple lines of evidence. The design and sequence information of PVC-loaded proteins, RNAs, and plasmids were provided, together with the preparation process of DNA/RNA and the primer sets employed in this work. The physicochemical characteristics of DART particles loaded with different RNAs were systematically evaluated through extracellular and intracellular fluorescence detection, fluorescence microscopy imaging, and quantitative analysis of delivery efficiency, accompanied by cytotoxicity assessments at different concentrations. High-throughput sequencing (NGS) was performed to verify gene editing outcomes, confirming the targeted knockout of GFP in A549 cells. Furthermore, structural insights into PVC tail fiber binding domains and their engineered variants were presented, and receptor-targeting models across multiple species were constructed, elucidating the cross-species adaptability of the delivery system. Together, these supplementary experiments and analyses establish a rigorous and coherent chain of validation for the DART platform, reinforcing the robustness and translational potential of our findings.

### Supplementary Figures

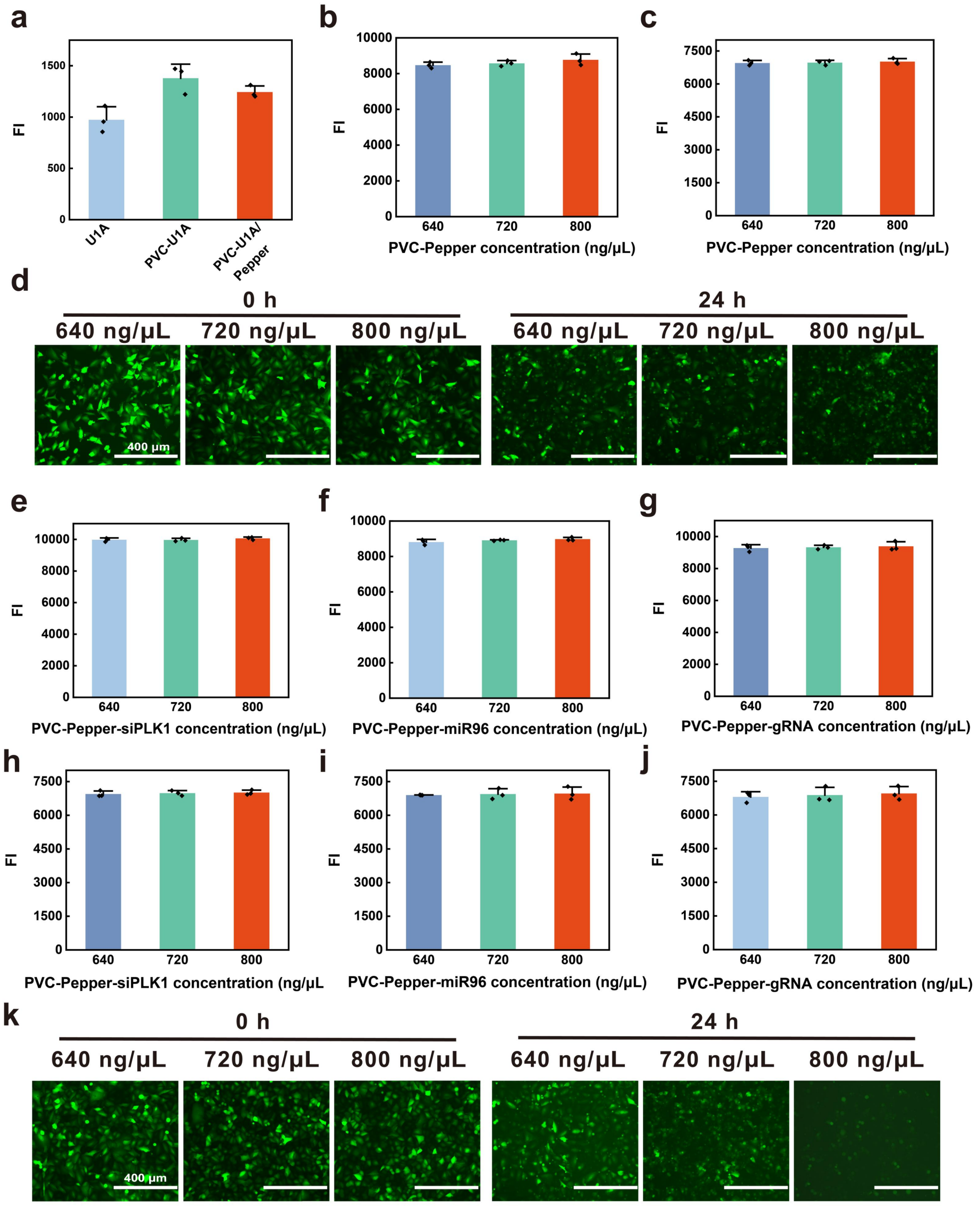


**Supplementary Figure 1 Fluorescence of DARTs loaded with different RNAs. a** Control experiment to detect whether Pepper was loaded into DART. **b–c** Extracellular (b) and intracellular (c) fluorescence detection results of DART loaded with Pepper RNA at 640, 720, and 800 ng/μL. **d** Fluorescence images of DART treating A549, indicating DART with high concentration was not suitable for working concentration. **e–g** Extracellular fluorescence detection results of DART loaded with Pepper- siPLK1 (e), Pepper-miR96 (f), and Pepper-gRNA (g) at 640, 720, and 800 ng/μL, respectively. **h–j** Intracellular fluorescence detection results of DART loaded with Pepper-siPLK1 (h), Pepper-miR96 (i), and Pepper-gRNA (j) at 640, 720, and 800 ng/μL, respectively. **k** Fluorescence images of DART treating A549, indicating DART with high concentration was not suitable for working concentration.

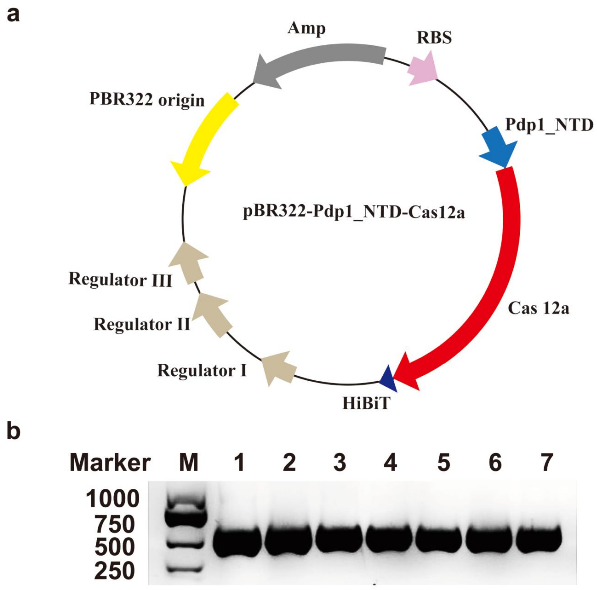


**Supplementary Figure 2 The construction of pBR322-Pdp1_NTD-Cas12a**. **a** Plasmid map of pBR322-Pdp1_NTD-Cas12a. **b** Agarose gel electrophoresis results of pBR322-Pdp1_NTD-Cas12a. Lane M: DNA marker (2000 bp, reference band indicated). Lanes 1–7: colony PCR products amplified from individual clones.

a

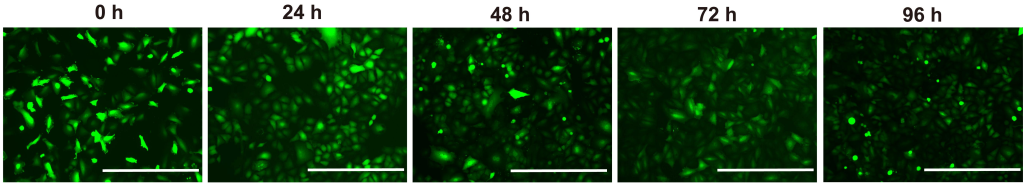


b

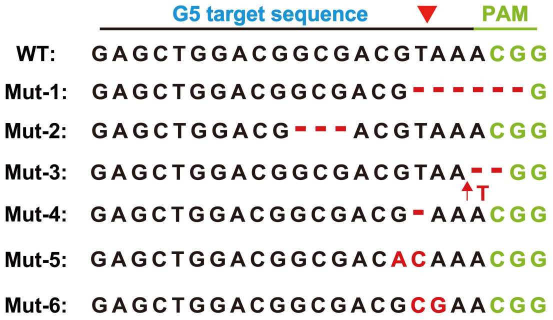


**Supplementary Figure 3 TatU1A mediated the knockout of GFP gene in vitro. a** Fluorescence microscopy of A549-GFP cells treated with TatU1A delivery for gRNA-G5 and PVC delivery for Cas9. **b** High-throughput sequencing of target loci revealed indel mutations after optimization of intracellular knockout experiments, including deletions, insertions, and substitutions following intracellular delivery of Cas9-G5 via PVC and TatU1A.

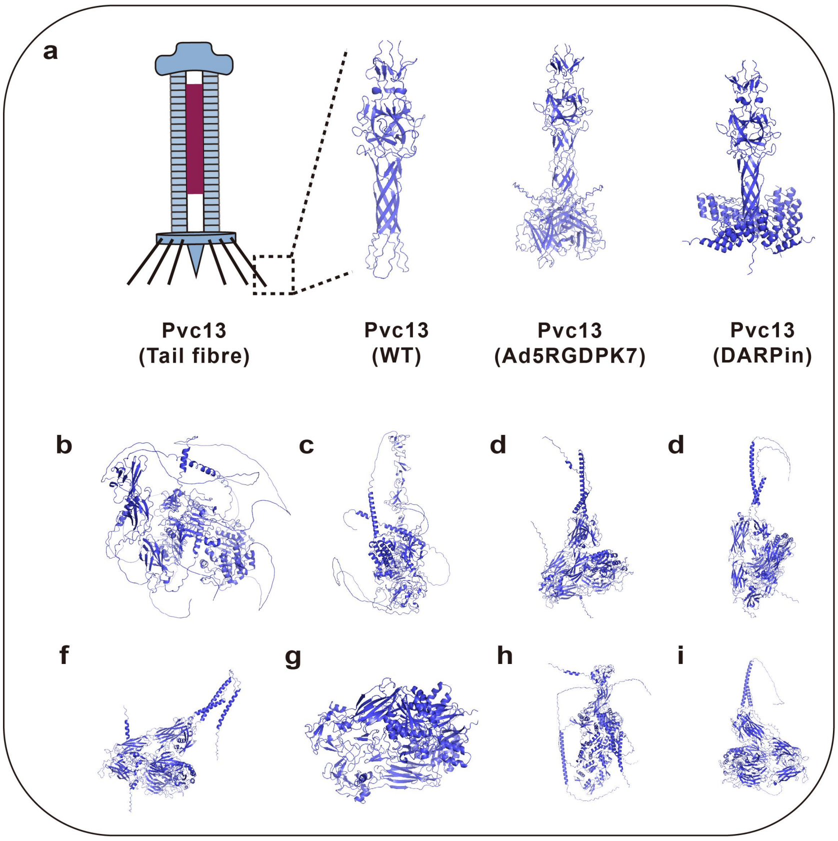


**Supplementary Figure 4 PVC tail fiber microstructure and receptor-targeting models. a** PVC13 wild-type binding domain (Pvc13 (WT)); the wild-type binding domain of Pvc13 was replaced with an extended tropism variant of adenovirus type 5 (Ad5) fiber apical sphere binding domain (Pvc13-Ad5RGDPK7) and an extended tropism variant of ankyrin repeat protein (DARPin) targeting epidermal growth factor receptor (EGFR) (Pvc13-DARPin). **b** Nematode growth factor receptor model. **c** Drosophila growth factor receptor model. **d** Zebrafish αVβ integrin receptor model. **e** mouse αVβ integrin receptor model. **f** Naked mole rat αVβ integrin receptor model. **g** pig αVβ integrin receptor model. **h** monkey αVβ integrin receptor model. **i** Human αVβ integrin receptor model.

#### Supplementary Tables

##### Supplementary Table 1: Design of PVC-loaded proteins and RNAs sequences

(a) The U1A protein was loaded into the PVC-loaded nucleic acid sequence by the Pd-NTD guidance sequence.

Pd_NTD+2xGGSGG linker+U1A+HiBiT

ATGCCTAGATATGCTAATTATCAGATAAACCCCAAACAGAATATTAAAAATTCACATGGG
AAATCTTCATCGTCAGATTTTTCTAGTGGGTACCTTTCATTCTCAAATAATTCGCTTGAT
GACCCTTTTATTCGGCAGCAAGTTAAGAGAGAATTTATTTGGGAGGGACATATGAAGG
AGATTGAAGAAGCTTCAAGATTAGGTGGCTCTGGTGGTGGCGGTTCTGGCGGC**GCAG
TTCCCGAGACCCGCCCTAACCACACTATTTATATCAACAACCTCAATGAGAAGATC
AAGAAGGATGAGCTAAAAAAGTCCCTGTACGCCATCTTCTCCCAGTTTGGCCAGA
TCCTGGATATCCTGGTATCACGGAGCCTGAAGATGAGGGGCCAGGCCTTTGTCAT
CTTCAAGGAGGTCAGCAGCGCCACCAACGCCCTGCGCTCCATGCAGGGTTTCCC
TTTCTATGACAAACCTATGCGTATCCAGTATGCCAAGACCGACTCAGATATCATTG
CCAAGATGAAATTA**GGGAGCTCCGGTGTGAGCGGCTGGCGGCTGTTCAAGAAGATTA
GC

(b) Nucleic acid sequence design of DART-loaded Pepper RNA anchored by U1A

T7 promoter+**UIA binding sequence**+linker1+**Pepper**+rrnB T1 terminator

TAATACGACTCACTATAGGG**GGGCATTGCACTCCGCCC**TCT**CGCACTGGCGCCGAG
GGCTTCCGCCAATCGTGGCGTGTCGGCG**AAATAAAACGAAAGGCTCAGTCGAAAG
ACTGGGCCTTTCGTTTTATCTGTTGTTTGTCGGTGAACGCTCTC

(c) Nucleic acid sequence design of DART-loaded gRNA anchored by U1A

T7 promoter+**UIA binding sequence**+linker1+**Pepper**+linker2+**gRNA**+rrnB T1 terminator

TAATACGACTCACTATAGGG**GGGCATTGCACTCCGCCC**TCT**CGCACTGGCGCCGAG
GGCTTCCGCCAATCGTGGCGTGTCGGCG**AGATCTT**AATTTCTACTCTTGTAGATGC
TTCTGTCCTAACAGGCTAG**AAATAAAACGAAAGGCTCAGTCGAAAGACTGGGCCTT
TCGTTTTATCTGTTGTTTGTCGGTGAACGCTCTC

(d) Nucleic acid sequence design of DART-loaded siPLK1 RNA anchored by U1A

T7 promoter+**UIA binding sequence**+linker1+**Pepper**+linker2+**siPLK1**+rrnB T1 terminator

TAATACGACTCACTATAGGG**GGGCATTGCACTCCGCCC**TCT**CGCACTGGCGCCGAG**
**GGCTTCCGCCAATCGTGGCGTGTCGGCG**AGATCTT**CAACCAAAGTGGAATATGAT**
**TGGGAAGTCATATTCCACTTTGGTTGTT**AAATAAAACGAAAGGCTCAGTCGAAAGA
CTGGGCCTTTCGTTTTATCTGTTGTTTGTCGGTGAACGCTCTC

(e) Nucleic acid sequence design of DART-loaded miR96 RNA anchored by U1A

T7 promoter+**UIA binding sequence**+linker1+**Pepper**+linker2+**miR96**+rrnB T1 terminator

TAATACGACTCACTATAGGG**GGGCATTGCACTCCGCCC**TCT**CGCACTGGCGCCGAG**
**GGCTTCCGCCAATCGTGGCGTGTCGGCG**AGATCTT**TGGCCGATTTTGGCACTAGC**
**ACATTTTTGCTTGTGTCTCTCCGCTCTGAGCAATCATGTGCAGTGCCAATATGGGA**
**AA**AAATAAAACGAAAGGCTCAGTCGAAAGACTGGGCCTTTCGTTTTATCTGTTGTTTG
TCGGTGAACGCTCTC

(f) Nucleic acid sequence design of DART-loaded gRNA-G5 anchored by U1A

T7 promoter+**gRNA-G5**+linker+**UIA binding sequence**+rrnB T1 terminator

TAATACGACTCACTATAGGG**GAGCTGGACGGCGACGTAAAGTTTTAGAGCTAGAAA**
**TAGCAAGTTAAAATAAGGCTAGTCCGTTATCAACTTGAAAAAGTGGCACCGAGTC**
**GGTGC**TCT**GGGCATTGCACTCCGCCC**CAAATAAAACGAAAGGCTCAGTCGAAAGAC
TGGGCCTTTCGTTTTATCTGTTGTTTGTCGGTGAACGCTCTC

(g) Nucleic acid sequence design of DART-loaded gRNA-G7 anchored by U1A

T7 promoter+**gRNA-G7**+linker1+**UIA binding sequence**+rrnB T1 terminator

TAATACGACTCACTATAGGG**CAGGGTCAGCTTGCCGTAGGGTTTTAGAGCUAGAAA**
**TAGCAAGTTAAAATAAGGCTAGTCCGTTATCAACTTGAAAAAGTGGCACCGAGTC**
**GGTGC**TCT**GGGCATTGCACTCCGCCC**CAAATAAAACGAAAGGCTCAGTCGAAAGAC
TGGGCCTTTCGTTTTATCTGTTGTTTGTCGGTGAACGCTCTC

(h) Nucleic acid sequence design of DART-loaded crRNA-G1 anchored by U1A

T7 promoter+**crRNA-G1**+linker+**UIA binding sequence**+rrnB T1 terminator

TAATACGACTCACTATAGGG**GGTTCCCAAGGACCTATATGGTTTTAGAGCTAGAAAT**
**AGCAAGTTAAAATAAGGCTAGTCCGTTATCAACTTGAAAAAGTGGCACCGAGTCG**
**GTGC**TCT**GGGCATTGCACTCCGCCC**CAAATAAAACGAAAGGCTCAGTCGAAAGACT
GGGCCTTTCGTTTTATCTGTTGTTTGTCGGTGAACGCTCT

**Supplementary Table 2 Synthesis and preparation of DNA or RNA**

| Name | DNA or RNA sequence |
|---|---|
| Pd_NTD | ATGCCTAGATATGCTAATTATCAGATAAACCCCAAACAGAATATTAAAAATTCACATGGAAATCTTCATC GTCAGATTTTTCTAGTGGGTACCTTTCATTCTCAAATAATTCGCTTGATGACCCTTTTATTCGGCAGCAAG TTAAGAGAGAATTTATTTGGGAGGGACATATGAAGGAGATTGAAGAAGCTTCAAGATTA |
| U1A binding sequence | GGGCAUUGCACUCCGCCC |
| Pepper RNA | CGCACUGGCGCCGAGGGCUUCCGCCAAUCGUGGCGUGUCGGCG |
| siPLK1 RNA | CAACCAAAGUGGAAUAUGAUUGGGAAGUCAUAUUCCACUUUGGUUGUU |
| miR96 RNA | UGGCCGAUUUUGGCACUAGCACAUUUUUGCUUGUGUCUCUCCGCUCUGAGCAAUCAUGUGCAGUG CCAAUAUGGGAAA |
| gRNA | AAUUUCUACUCUUGUAGAUGCUUCUGUCCUAACAGGCUAG |
| gRNA-G5 | GAGCUGGACGGCGACGUAAAGUUUUAGAGCUAGAAAUAGCAAGUUAAAAUAAGGCUAGUCCGUUA UCAACUUGAAAAAGUGGCACCGAGUCGGUGC |
| gRNA-G7 | CAGGGUCAGCUUGCCGUAGGGUUUUAGAGCUAGAAAUAGCAAGUUAAAAUAAGGCUAGUCCGUUA UCAACUUGAAAAAGUGGCACCGAGUCGGUGC |
| crRNA-G1 | AAUUUCUACUCUUGUAGAUCGUCGCCGUCCAGCUCGACCAGG |

#### Supplementary Table 3 Prime sequence

| Primer name | Prime sequence |
|---|---|
| AR-oriLT-PcrR0107 | agaatccaagcactagtcatgac |
| UIApep-F-0510 | gcgccgagggcttccgccaatcgtggcgtgtcggcgagatcttaatttctactcttgtagatgcttctg |
| AF-tAR-PR0107 | gtcatgactagtgcttggattct |
| UIApep-R-0510 | ttggcggaagccctcggcgccagtgcgagagggcggagtgcaatgcccttctctatcactgatagggag |
| UPR-Y-F | ccttcgattccgacctcatt |
| UPR-Y-R | cctttgagtgagctgatacc |
| U1A-Amp-F | cacgttaagggattttggtc |
| U1A-Amp-R | gaactgcactagtgccgccagaaccgccaccac |
| U1A-ori-F1 | gatgaaattagggagctccggtgtgagcgg |
| U1A-ori-R | gaccaaaatcccttaacgtg |
| U1A-p-F | tggcggcactagtgcagttcccgagacccgccc |
| U1A-p-R | cggagctccctaatttcatcttggcaatgatatctgagt |
| UIA-AMP-YF1 | gcttagaaagactcggtgcc |
| UIA-AMP-YR1 | ttcaggctccgtgataccag |
| UIA-ORI-YF1 | ctatgcgtatccagtatgcc |
| UIA-ORI-YR1 | agtgaaactaaccttcctcc |
| U1A-lys-R | cccctatagtgagtcgtattaatctcgagtgataaactctccgtgatgg |
| U1A-ori-F2 | ctagggagacgtcgaccgatgcccttgaga |
| U1A-pep-F | agattaatacgactcactatagggggggcattgcactccgccctc |
| U1A-pep-R | atcggtcgacgtctccctaggtataaacgc |
| OU-YF1 | gagtcacactggctcacctt |
| OU-YR1 | gaatcataatggggaaggcc |
| ZU-YF2 | gtgtctctcatcctgaagaa |
| ZU-YR2 | tagaaattaagatctcgccg |
| PLK1-R-0326 | cgactgagcctttcgttttatttaacaaccaaagtggaatatgacttcccaatcatattccactttggttgaagatctcgccgacacgc |
| RT-F-0326 | aaataaaacgaaaggctcagtcg |
| PLK1-YF | ggaatatgattgggaagtca |
| G1-YR-0821 | taatggggaaggccatccag |
| PLK1-YR | gtggaatatgacttcccaat |
| CreT-F-0326 | aagatgatactctggctggcatctgtccttggaaacactcatgccagccacgatcaggaaataaaacgaaaggctcagtcg |
| CreT-R-0326 | tccaaggacagatgccagccagagtatcatcttctcatgtgatcacccttcttaagatctcgccgacacgcc |
| CreT-YF | ggtgatcacatgagaagatg |
| CreT-YR | agtgtttccaaggacagatg |
| mi96-F-0326 | actagcacatttttgcttgtgtctctccgctctgagcaatcatgtgcagtgccaatatgggaaaaaataaaacgaaaggctcagtcg |
| mi96-R-0326 | cacaagcaaaaatgtgctagtgccaaaatcggccaaagatctcgccgacacgcca |
| miR96-YF | ggcactagcacatttttgct |
| miR96-YR | tgtgctagtgccaaaatcgg |
| RT-F-0326 | aaataaaacgaaaggctcagtcg |
| TR-R-0326 | cgactgagcctttcgttttatttcgcactaaggcaatgggtttcagggcttgcatggctaagatctcgccgacacgcca |
| TTR-YF | tagccatgcaagccctgaaa |
| TRR-YR | cactaaggcaatgggtttca |
| ArcZ-F-0410 | tggtgttggcgcagtattcgcgcaccccggtctagccggggtcatttttaaataaaacgaaaggctcagtcg |
| ArcZ-R-0411 | cgaatactgcgccaacaccagggaaaaagatctcgccgacacgcca |

| | |
|---|---|
| ArcZ-YF | tggtgttggcgcagtattcg |
| ArcZ-YR | aaaatgaccccggctagacc |
| C12-F-pd2-0821 | ggtctgacagttaccaatgcttaatcagtg |
| C12-R-pd2-0821 | tcttcagcattgctgccggaacaccatgaa |
| C12-F-pd1-0821 | agcggctggcggctgttcaagaagattagctgatatggattttcatgtcatcaggagaa |
| C12-R-pd1-0821 | gcattggtaactgtcagaccaagtttactc |
| C12-F-cas-0821 | tccggcagcaatgctgaagaacgtgggcat |
| C12-R-cas-0821 | cttcttgaacagccgccagccgctcacaccggagctcccgtgtttcacgctggtctgag |
| C12-Y1F-0821 | gccttctacagcagcttcat |
| C12-Y1R-0821 | cacaggcccatagttccagg |
| C12-Y2F-0821 | aagactcggtgccagtgtaa |
| C12-Y2R-0821 | tctcaggttgatctccaggt |
| G1-F1-0821 | actcttgtagatcgtcgccgtccagctcgaccaggcaaataaaacgaaaggctcagtcg |
| G1-R2-0821 | tcgagctggacggcgacgatctacaagagtagaaattaaagagggcggagtgcaatgcc |
| G1-YF-0821 | cgtcgccgtccagctcgacc |
| G1-YR-0821 | taatggggaaggccatccag |
| G4-F1-0821 | ctagtccgttatcaacttgaaaaagtggcaccgagtcggtgctctgggcattgcactccgccctcaaataaaa cgaaaggctcagtcga |
| G4-R2-0821 | gccactttttcaagttgataacggactagccttattttaacttgctatttctagctctaaaacccctatagtgagtcgtattaatctcg |
| G4-YF-0821 | taaggctagtccgttatcaacttgaa |
| G5-F1-0821 | gagctggacggcgacgtaaagttttagagctagaaatagcaagttaaaat |
| G5-R2-0821 | tttacgtcgccgtccagctcccctatagtgagtcgtattaatctcg |
| G5-YF-0821 | gagctggacggcgacgtaaa |
| G7-F1-0821 | cagggtcagcttgccgtagggttttagagctagaaatagcaagttaaaat |
| G7-R2-0821 | cctacggcaagctgaccctgccctatagtgagtcgtattaatctcg |
| G7-YF-0821 | cagggtcagcttgccgtagg |

**Supplementary Table 4 Strains and plasmids used in this study.** Note: Data presented under "Origin" column; "Our laboratory" denotes constructs generated in this work, "Addgene" indicates commercial plasmids (catalog numbers provided in Methods).

| Strain/Plasmid/Primer | Characteristics | Origin |
|---|---|---|
| *E. coli* BL21 (DE3) | Protein expression | This study |
| *E. coli* DH5α | Plasmid construction | This study |
| *E. coli* EPI300 | PVC expression | This study |
| pBR322-Pdp1_NTD-GFP-HiBiT | For assembling PVC payload modules[4] | Addgene |
| pET28a-Tat-U1A | used as the pBR322-UIA amplification template | This study |
| pBR322-UIA | For loading U1A into PVCs | This study |
| pBR322-UIA-Pepper | For loading U1A/pepper RNA complex into PVCs | This study |
| pBR322-U1A-gRNA | For loading U1A/RNA complexes into PVCs | This study |
| pBR322-U1A-siPLK1 | For loading U1A/RNA complexes into PVCs | This study |
| pBR322-U1A-miR96 | For loading U1A/RNA complexes into PVCs | This study |
| pAWP78-PVCpnf_pvc13-Ad5RGDPK7 | For expressing PVC structural machinery[1] | Addgene |
| pAWP78-PVCpnf_pvc13-E01DARPin | For expressing PVC structural machinery[2] | Addgene |
| pAWP78-PVCpnf_pvc13-antiMouseMHCIINb | For expressing PVC structural machinery[3] | Addgene |
| pBR322-Pdp1_NTD-ZFD_L | For assembling PVC payload modules[5] | Addgene |
| pET28TEV-LbCpf1 | For assembling PVC payload modules | This study |
| pBR322-Pdp1_NTD-Cas9 | For loading Cas9 into PVCs[4] | Addgene |
| pBR322-Pdp1_NTD-Cas12a | For loading Cas12a into PVCs | This study |
| pBR322-Pd_U1A-crRNA-G1 | For loading sgRNA modules into PVCs | This study |
| pBR322-Pd_U1A-gRNA-G5 | For loading sgRNA modules into PVCs | This study |
| pBR322-Pd_U1A-gRNA-G7 | For loading sgRNA modules into PVCs | This study |

**Supplementary Table 5 Cell lines used in this study.**

| *Cell line* | *Organism* | *Source* | *Media* |
|---|---|---|---|
| A549 | Human (H. sapiens) | ATCC CCL-185 | RPMI+GlutaMAX |

**Supplementary Table 6 Comparison of DART with existing delivery vectors**

| Vector Type | Advantages | Disadvantages / Notes | Market Price | Sources |
|---|---|---|---|---|
| Adeno-Associated Virus (AAV) | Low immunogenicity; enables long-term expression in both dividing and non-dividing cells; FDA-approved for multiple gene therapy products | Limited cargo capacity (~4.7 kb); complex and costly production | ~$3,000–$5,000 per research-grade batch; commercial gene therapy products priced >$1M per dose | https://www.vectorbiolabs.com/ |
| Lentivirus | Stable genome integration; suitable for long-term expression; high transduction efficiency in diverse cell types | Potential risk of insertional mutagenesis; complex production | ~$1,500–$3,000 per research-grade batch; key cost driver in CAR-T therapies | https://www.addgene.org/ |
| Adenovirus | High transduction efficiency; large cargo capacity (up to 30–36 kb); scalable production | Strong immunogenicity; short duration of in vivo expression; requires high doses | ~$1,000–$2,000 per research-grade batch; vaccines (e.g., AstraZeneca COVID-19) cost ~$3–$5 per dose | https://www.atcc.org/ |
| Herpes Simplex Virus (HSV) | Very large cargo capacity (>30 kb); neuronal targeting; applicable in neurological disorders | Potential toxicity; complex regulation; long-term safety concerns | commercial oncolytic virus (T-VEC, Imlygic) priced >$65,000 per treatment | https://www.imlygic.com/ |
| Retrovirus (γ-retrovirus) | Stable genome integration; historically used in early clinical gene therapy | High risk of insertional mutagenesis; only transduces dividing cells | ~$1,000–$2,000 per research-grade batch | https://www.addgene.org/ |
| Lipid Nanoparticles (LNP) | High encapsulation efficiency; clinically validated in mRNA vaccines | Stability and storage issues; potential immunogenicity | ~$ 20 (per COVID-19 vaccine dose) | https://www.ema.europa.eu |
| Polyethylenimine (PEI) | High transfection efficiency; inexpensive | Cytotoxicity; low biocompatibility | ~$ 200 (per research reagent kit, not clinical drug) | https://www.sigmaaldrich.com |
| Poly(lactic-co-glycolic acid) (PLGA) | Biodegradable; FDA-approved material | Limited nucleic acid delivery efficiency | ~$ 150 (per research kit, not marketed drug) | https://www.fda.gov |
| Exosome-based Delivery | Natural biocompatibility; potential for precision medicine | Low yield; scalability issues | ~$ 1,000–2,000 (per experimental dose in clinical trials) | https://clinicaltrials.gov |
| GalNAc Conjugates | High hepatocyte specificity; subcutaneous administration; clinically proven | Limited to liver-targeted diseases | ~$319.92 per dose (Inclisiran, NHS UK price) | https://www.nice.org.uk |
| DART | Biodegradable, non-immunogenic, cost-effective, simple to produce, capable of delivering and protecting RNA | Puncture-Induced Membrane Damage | Each experiment employed a 1 mL (5mg $mL^{-1}$) dosage, incurring a cost of approximately ~$2.38 | This study |

**Supplementary Table 7 Cost calculation of DART (per 1 mL)**

| Category | Component | Amount | Unit Price (USD) | Cost (USD) |
|---|---|---|---|---|
| Lysis Buffer (1 L) | Tris | 121.1400 g | 0.0534/g | 6.4800 |
| | NaCl | 8.1820 g | 0.0014/g | 0.0113 |
| | KCl | 0.2240 g | 0.0251/g | 0.0056 |
| | $MgCl_2 \cdot 6H_2O$ | 1.0170 g | 0.0019/g | 0.0019 |
| | Triton X-100 | 5.0000 mL | 0.0836/mL | 0.4181 |
| Total (1 L) | - | - | - | 6.9169 |
| For 28 mL | - | - | - | 0.1936 |
| LB Medium (1 L) | Tryptone | 10.0000 g | 0.0362/g | 0.3625 |
| | Yeast extract | 5.0000 g | 0.0669/ g | 0.3345 |
| | NaCl | 10.0000 g | 0.0014/g | 0.0138 |
| Total (1 L) | - | - | - | 0.7108 |
| For 0.5 L | - | - | - | 0.3554 |
| Enzymes & Inhibitors | Protease inhibitor | 0.2900 mL | 4.1800/mL | 1.2122 |
| | DNase | 0.0560 mL | 7.8000/mL | 0.4373 |
| | Lysozyme | 0.0560 mL | 3.2500/ mL | 0.1825 |
| Total | - | - | - | 1.8320 |
| Overall Cost (1 mL DART) | - | - | - | 2.3810 |

### Supplementary Methods

**Predicting Protein Complex Structures Using AlphaFold 3.0**

AlphaFold 3.0 (DeepMind, v3.0.1) was used to predict the structure of cross-species receptor complexes. Firstly, FASTA sequences of target receptors were obtained from UniProt, including nematodes, fruit flies, zebrafish, mice, rats, pigs, monkeys, and humans. The task is submitted through Alphafold Server (https://alphafoldserver.com/), and the key parameter settings are : num _ recycles = 12 (enhanced multi-chain convergence), num _ ensemble = 8 (conformational space sampling). The output model was double filtered by pLDDT⩾70 and interface PAE<5Å, and the optimal structure was visualized by PyMOL 2.5.

**CRISPResso2-based quantification of genome editing efficiency**

Based on the Illumina double-end sequencing data, the adaptor was first removed by Cutadapt (v4.4) and quality control (Phred ≥ 30), and then CRISPResso2 (v2.3) was used for core analysis (parameters: amplicon sequence 200–300 bp, gRNA flanking window 10 bp, minimum alignment similarity 85%, single base accuracy detection). After quality control screening with depth ≥ 200× and effective reading ≥ 10,000, bilateral Fisher test was used to quantify editing efficiency (FDR corrected q99 % consistency) and negative and positive controls[6].

**In vivo DART clearance assay**

To evaluate the persistence of DART in subcutaneous tissue of mice, we isolated and analyzed the skin tissue fluid at the injection site[4]. The specific steps were as follows:

1. Tissue collection and processing: After euthanasia of DART-treated mice, the skin tissue at the injection site was dissected and separated using a sterile scalpel. The obtained tissues were rinsed with PBS for three consecutive times.
2. Homogenization and primary centrifugation: The tissue was prepared into a suspension using a Dounce homogenizer. The suspension was then centrifuged at 500 ×g for 5 min.
3. Supernatant treatment and DART enrichment: Clarified supernatant after centrifugation was collected, diluted with PBS to 28 mL, and centrifuged at 120,000 ×g at 4°C for 2 h. The aim of this step WAS to precipitate the complete DART complex that might exist.
4. Washing of sediment: The sediment obtained by ultracentrifugation was resuspended in 50 μL PBS and centrifuged at 16,000 ×g for 15 min at 4°C to remove residual crude homogenate impurities.
5. DART complex detection: Finally, the re-suspended precipitate after washing was used for negative staining transmission electron microscopy ( negative staining TEM ) analysis to detect the complete DART complex (the specific method is detailed in the manuscript 'Electron microscope' section).
